# Adaptation of STARR-seq method to be used with 3^rd^ generation integrase-deficient promoterless lentiviral vectors

**DOI:** 10.1101/2020.06.24.169714

**Authors:** Mantas Matjusaitis, Donato Tedesco, Azuolas Ciukas

## Abstract

Ability to functionally screen gene regulatory sequences, such as promoters and enhancers, in high throughput manner is an important prerequisite for many basic and translational research programs. One of the methods that allow such screening is STARR-seq, or self-transcribing active regulatory region sequencing. It allows to quickly screen millions of candidate sequences in the cell type of interest. However, it does rely on transfection as a delivery method which can be a limiting-step for some hard-to-transfect cells such as senescent cells. Here we show that integration-deficient and integration-competent promoterless lentiviral particles can be used to deliver STARR-seq constructs into cells. These constructs reported CMV enhancer activity both at protein and mRNA level. While further validations are necessary, ability to deliver STARR-seq libraries using lentiviral particles will significantly improve the versatility and usability of such a method.

## INTRODUCTION

Discovery of novel endogenous promoters, enhancers and other gene regulatory sequences plays a critical role in a number of research fields. These elements contain binding motifs for various transcription factors (TFs), co-factors and mediators. As such, they act as hubs from which transcription can be initiated, enhanced and/or repressed. In general terms, promoters are typically located close to genes it regulates, while enhancers are more distal elements often operating tens and even hundreds of kilobases away from gene coding region (Sanyal et al., 2012). Furthermore, it is estimated that these cis-regulatory elements significantly outnumber genes (Shen et al., 2012). While due to the nature of these sequences it can be difficult to identify and characterize them, it is an important task. Such research can not only help to depict how biological systems work but also facilitate invention of new synthetic promoters that can be used in gene therapies to achieve cell and tissue specificity (Piekarowicz et al., 2019).

One prominent way of scoring enhancer activity is known as self-transcribing active regulatory region sequencing or STARR-seq (Arnold et al., 2013). In this method, suspected enhancer sequences are cloned between minimal promoter (followed by an open reading frame of a reporter) and polyadenylation (polyA) signal. Once such a vector is delivered into cells by transfection, it will transcribe from the core promoter at a very low level. If enhancer candidate sequence acts as a functional enhancer it will increase transcription resulting in a higher abundance of that particular mRNA species. Because enhancer candidate sequence is located before polyA signal, its sequence will be included in the mRNA. With the help of sequencing, most abundant mRNA species - and therefore strongest enhancers - can be identified (Fig.1A). In recent years, this method was further developed by combining it with ATAC, ChIP and solving some of the systemic problems giving raise to false-positives (Muerdter et al., 2018; Vockley et al., 2016; Wang et al., 2018). While it is a powerful way of scoring candidate enhancers, it can be difficult to transfect some cell lines, for example, senescent cells. A popular alternative to transfection in hard-to-transfect cells is viral transduction - delivery of nucleic acid into cells using viral particles.

**Figure 1.**
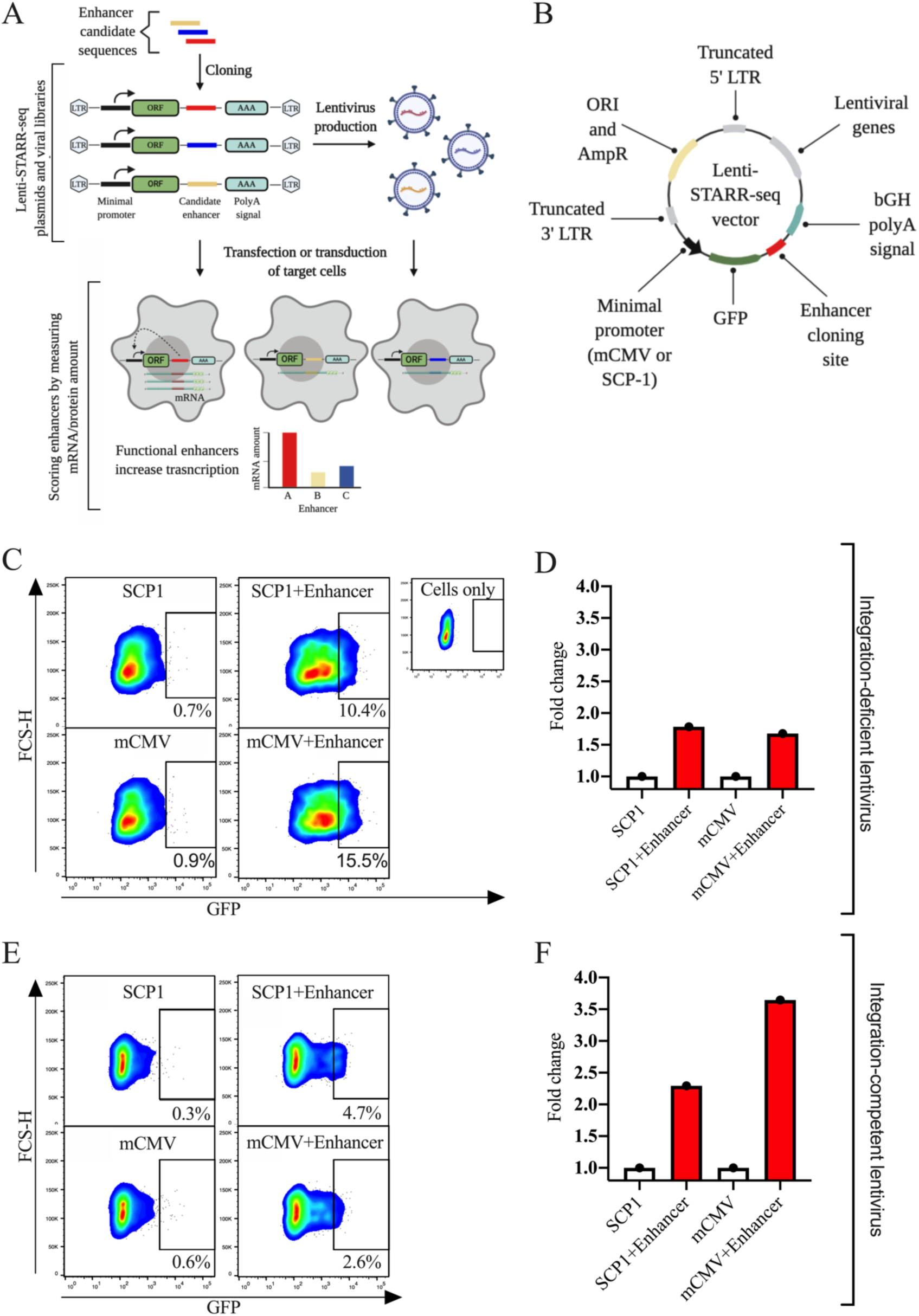
Adoption for STARR-seq method to be used with lentiviral vectors. (**A**) Graphical representation of the process for making Lenti-STARR-seq libraries. Candidate enhancer sequences are cloned into Lenti-STARR-seq plasmids. These plasmids can then either be used to transfect cells or produce lentivirus particles. Transfected or transduced cells are analysed by measuring reporter and/or mRNA levels to identify which enhancers have positively affected transcription. (**B**) Structure of Lenti-STARR-seq vector. (**C**) Flow cytometry and (**D**) RT-qPCR data from U2OS cells 48 hours after transduction with integration-deficient Lenti-STARR-seq vectors with (“SCP1+Enhancer” and “mCMV+Enhancer”) or without (“SCP1” and “mCMV”) CMV enhancer, n=1. (**E**) Flow cytometry and (**F**) RT-qPCR data from U2OS cells 48 hours after transduction with integration-competent Lenti-STARR-seq vectors with (“SCP1+Enhancer” and “mCMV+Enhancer”) or without (“SCP1” and “mCMV”) CMV enhancer, n=1.

A number of different viruses have been adapted for safe use in research. Arguably, most common of them being lentiviral vectors. Lentivirus is a subtype of retrovirus that is able to infect both dividing and non-dividing cells. Upon infection, lentivirus integrates into a host genome – generally into actively transcribed genes (Papanikolaou et al., 2015). In such a fashion, it can be used as a tool to easily deliver diverse libraries of payloads (up to 8kb in size) into a large number of cells, both for clinical and research purposes. In this study, we report proof of principle data demonstrating how STARR-Seq plasmids could be adapted for use with integration-competent and integration-deficient promoterless lentiviral systems.

## RESULTS

### Design and construction of Lenti-STARR-seq plasmids

To construct a lentiviral vector which could be used together with STARR-seq method, a number of considerations and modifications were necessary (Fig.1B, Fig.S1 for full sequence). Firstly, 5’ and 3’ LTR sequences have been shown to work as promoter/enhancers (Klaver & Berkhout, 1994; Medstrand et al., 2001). This would likely interfere with the signal-to-noise ratio of an assay, therefore promoterless lentiviral vectors (lacking U3 regions in both LTRs; also known as self-inactivating or SIN) will be used (Yu et al., 1986). Secondly, to avoid splicing out an intron - which is an element of STARR-seq plasmid - and to minimize any remaining interference from Δ5’LTR, a whole construct will be placed in a reverse orientation (Cooper et al., 2015; Poling et al., 2017). This should also help to lower risks of sub-optimal viral production due to the presence of bGH polyA sequence between LTRs (Fig. 1B) (Hager et al., 2008). Thirdly, while the integration of STARR-seq screening vectors into genome did not interfere with signal in previous studies (Arnold et al., 2013), lentiviral integration into actively transcribed genome loci could result in false positives or higher noise. To mitigate this risk, we will investigate if integration-deficient particles can be used (Wanisch & Yáñez-Muñoz, 2009). Integration-deficient viral genomes form DNA episomes that can persist for weeks and can be harvested using commercially available kits – which would be important for testing library delivery and coverage during larger screen (Badralmaa & Natarajan, 2013; Vargas et al., 2004). Lastly, we have constructed two Lenti-STARR-seq versions containing two different minimal promoters – mCMV and SCP1. While our plasmids also contain bacterial ORI - which has been shown to function as an enhancer and enhance noise (Muerdter et al., 2018) – it will not be packaged into lentiviral particles. mTagGFP reporter was placed downstream of these minimal promoters to allow for an alternative method of screening for strong enhancers (Fig.1B). Constructed plasmids were verified using sequencing.

### Validation of Lenti-STARR-seq vectors

To test if Lenti-STARR-seq lentiviral vectors can be used to score enhancer activity, CMV enhancers were cloned into both Lenti-STARR-seq plasmid variants (mCMV and SCP1) using Bpi restriction sites. After production of integration-deficient lentiviral particles, U2OS cells were transduced. 48 hours post-transduction cells were analysed by flow cytometry (Fig.1C) and RT-qPCR (Fig.1D). mTagGFP signal of vectors lacking CMV enhancer (“SCP1” and “mCMV”) was higher compared to non-transduced cells (“Cells only”) suggesting some basal transcription activity in these vectors. Importantly, the addition of CMV enhancer to these vectors (“SCP1+Enhancer” and “mCMV+Enhancer”) increase the amount of mTagGFP positive cells (Fig.1C). In addition to that, a similar trend can be observed at mRNA level (Fig.1D). Integration-competent lentivirus particles were also tested. Similarly to previous observations, an increase in signal both at the protein and at mRNA level was observed after addition of CMV enhancer (Fig.1E and Fig.1F).

## DISCUSSION

Ability to functionally screen candidate enhancer activity in a high-throughput assay is immensely useful for a number of research fields. One of such powerful approach is STARR-seq method which can be used to evaluate thousands of candidates with high sensitivity. However, one limitation of such method can be the inability to effetely delivery these DNA constructs into cells. In this short study, we present preliminary data that STARR-seq method can be used with an integrase-deficient lentiviral delivery system. This adoption can significantly enhance STARR-seq capabilities and usability.

A number of considerations have been taken into account to merge lentiviral delivery with STARR-seq constructs. This includes the use of integrase-deficient lentiviral packaging, reversing the orientation of STARR-seq construct and use of SIN promoterless LTRs. Preliminary data suggest that these vectors are capable of reporting activity of CMV enhancer both at protein and mRNA level. While signal between controls (constructs with only a minimal promoter) and samples (constructs with a minimal promoter and an enhancer) was evident, a level of noise was also observed then comparing non-transduced cells to cells containing a construct with a minimal promoter only. Further work might be necessary to optimise the signal-to-noise ratio.

While this data offers interesting insights into how STARR-seq method could be used with integration-deficient promoterless lentiviral vectors, a number of caveats and future work directions need to be mentioned. Firstly, because STARR-seq method relies on sequencing as a readout, it will be important to validate Lenti-STARR-seq approach in this context. Secondly, this study would be improved if viral production quality (including titres) and integration events (or lack of them) would be monitored. Lastly, while multiple independent transductions were performed, it is necessary to validate these findings with biological replicates to allow for statistical analysis. Taken together, we would like to highlight that this study is only preliminary and further work is needed to validate and optimize Lenti-STARR-seq vectors.

## MATERIALS AND METHODS

### Cloning

Lenti-STARR-seq plasmids, containing either mCMV or SCP-1 minimal enhancers, were constructed using a combination of gene synthesis and restriction enzyme cloning. The full sequence of Lenti-STARR-seq vectors can be found in supplementary data. For validating the functionality of Lenti-STARR-seq plasmids, CMV enhancer was cloned in using Bpi restriction sites.

mCMV sequence:

5’GTAGGCGTGTACGGTGGGAGGTCTATATAAGCAGAGCTCGTTTAGTGAACCGTCAGATC

SCP-1 sequence:

5’gtacttatataagggggtgggggcgcgttcgtcctcagtcgcgatcgaacactcgagccgagcagacgtgcctacggaccg

CMV enhancer sequence:

5’catggtaatagcgatgactaatacgtagatgtactgccaagtaggaaagtcccgtaaggtcatgtactgggcataatgccaggcgggccatttaccgtcattgacgtcaatagggggcggacttggcatatgatacacttgatgtactgccaagtgggcagtttaccgtaaatactccacccattgacgtcaatggaaagtccctattggcgttactatgggaacatacgtcattattgacgtcaatgggcgggggtcgttgggcggtcagccaggcgggccatttaccgtaagttatgtaacgcggaactccatatatgggctatgaactaatgaccccgtaattgattactattaataactagtcaataatcaatgcc

### Lentivirus production

Lentivirus was produced by transient transfection of Lenti-STARR-seq vector, vsv-g vector pMD2G (Addgene #12259) and pspax2 vector (Addgene #12260) or pspax2 vector carrying D64A mutant viral integrase (for integrase-deficient lentivirus production). Vectors were transfected at 2:1:2 ratio into HEK293T cells using lipofectamine (TheromFisher), using manufacturer’s recommendations. The supernatant containing virus was harvested and filtered through 0.2um PES filter 48h after transfection.

### Cell culture and transduction

U2OS cell line has been used in all of the experiments. Cell were routinely cultured in DMEM containing 10% FBS, without antibiotics.

Transductions were performed in 12-well plate format. Each well contained 1×105 cells with 1ml DMEM, 10% FBS and polybrene 5ug/ml. 300μl (integration-deficient virus) or 50μl (integration-component virus) of filtered virus supernatant was added to cells.

### RNA extraction, RT and RT-qPCR

48 hours after transduction, cells were harvested by trypsinization. Total RNA was extracted using RNeasy Mini Kit (Qiagen, #74104) according to the manufactures’ protocol. 400ng of total RNA was used to produce cDNA using MMLV High Performance Reverse Transcriptase (Lucigen RT80125K). TaqMan qPCR was performed using PrimeTime® Gene Expression Master Mix IDT; # 1055772). The following oligos targeting GFP and GAPDH were used:

TagGFP2 probe: CCTCGGGCATGGCGCTCTTG

TagGFP2_forward: TGGTGCGCTCCTGGATGTAG

TagGFP2_reverse: CCCGAGCACATGAAGATGAAC

GAPDH probe: CGTTCTCAGCCTTGACGGTGCCA

GAPDH forward: GATTCCACCCATGGCAAATTC

GAPDH reverse: GGGATTTCCATTGATGACAAGC

Each sample was run as technical triplicate. Fold change, relative to GAPDH, was calculated using DataAssist software (ThermoFisher).

### Flow cytometry

Cells were harvested and analysed on FACS Melody cell sorter (BD Biosciences). Unless otherwise stated, 3,000 events were collected in each experiment. mTagGFP fluorescence data was collected using 488 nm laser and 530/30nm bandpass filter. Data was analysed using FlowJo software.

### Data analysis

Data was processed using GraphPad Prism, FlowJo and DataAssist software. Illustrations were created with BioRender.com.

## ACKNOWLEDGEMENTS

This work was supported by Life Extension Advocacy Foundation (LEAF) and its community, Longecity Community, Mike Nelson and Alan Mole.

## AUTHOR CONTRIBUTIONS

M.M. conceived and supervised the project; M.M. and A.C. acquired the funding; M.M. and A.C. wrote and edited the manuscript; D.T. supervised and performed experiments.

## REFERENCES

Arnold, C. D., Gerlach, D., Stelzer, C., Boryn, L. M., Rath, M., & Stark, A. (2013). Genome-wide quantitative enhancer activity maps identified by STARR-seq. Science, 339(6123), 1074–1077. https://doi.org/10.1126/science.1232542

Badralmaa, Y., & Natarajan, V. (2013). Impact of the DNA extraction method on 2-LTR DNA circle recovery from HIV-1 infected cells. Journal of Virological Methods, 193(1), 184–189. https://doi.org/10.1016/j.jviromet.2013.06.014

Cooper, A. R., Lill, G. R., Gschweng, E. H., & Kohn, D. B. (2015). Rescue of splicing-mediated intron loss maximizes expression in lentiviral vectors containing the human ubiquitin C promoter. Nucleic Acids Research, 43(1), 682–690. https://doi.org/10.1093/nar/gku1312

Hager, S., Frame, F. M., Collins, A. T., Burns, J. E., & Maitland, N. J. (2008). An internal polyadenylation signal substantially increases expression levels of lentivirus-delivered transgenes but has the potential to reduce viral titer in a promoter-dependent manner. Human Gene Therapy, 19(8), 840–850. https://doi.org/10.1089/hum.2007.165

Klaver, B., & Berkhout, B. (1994). Comparison of 5’ and 3’ Long Terminal Repeat Promoter Function in Human Immunodeficiency Virus. J Virol., 68(6): 383. https://pubmed.ncbi.nlm.nih.gov/8189520/

Medstrand, P., Landry, J. R., & Mager, D. L. (2001). Long terminal repeats are used as alternative promoters for the endothelin B receptor and apolipoprotein C-I genes in humans. Journal of Biological Chemistry, 276(3), 1896–1903. https://doi.org/10.1074/jbc.M006557200

Muerdter, F., Boryn, Ł. M., Woodfin, A. R., Neumayr, C., Rath, M., Zabidi, M. A., Pagani, M., Haberle, V., Kazmar, T., Catarino, R. R., Schernhuber, K., Arnold, C. D., & Stark, A. (2018). Resolving systematic errors in widely used enhancer activity assays in human cells. Nature Methods, 15(2), 141–149. https://doi.org/10.1038/nmeth.4534

Papanikolaou, E., Paruzynski, A., Kasampalidis, I., Deichmann, A., Stamateris, E., Schmidt, M., Von Kalle, C., & Anagnou, N. P. (2015). Cell cycle status of CD34 + hemopoietic stem cells determines lentiviral integration in actively transcribed and development-related genes. Molecular Therapy, 23(4), 683–696. https://doi.org/10.1038/mt.2014.246

Piekarowicz, K., Bertrand, A. T., Azibani, F., Beuvin, M., Julien, L., Machowska, M., Bonne, G., & Rzepecki, R. (2019). A Muscle Hybrid Promoter as a Novel Tool for Gene Therapy. Molecular Therapy - Methods and Clinical Development, 15, 157–169. https://doi.org/10.1016/j.omtm.2019.09.001

Poling, B. C., Tsai, K., Kang, D., Ren, L., Kennedy, E. M., & Cullen, B. R. (2017). A lentiviral vector bearing a reverse intron demonstrates superior expression of both proteins and microRNAs. RNA Biology, 14(11), 1570–1579. https://doi.org/10.1080/15476286.2017.1334755

Sanyal, A., Lajoie, B. R., Jain, G., & Dekker, J. (2012). The long-range interaction landscape of gene promoters. Nature, 489(7414), 109–113. https://doi.org/10.1038/nature11279

Shen, Y., Yue, F., Mc Cleary, D. F., Ye, Z., Edsall, L., Kuan, S., Wagner, U., Dixon, J., Lee, L., Ren, B., & Lobanenkov, V. V. (2012). A map of the cis-regulatory sequences in the mouse genome. Nature, 488(7409), 116–120. https://doi.org/10.1038/nature11243

Vargas, J., Gusella, G. L., Najfeld, V., Klotman, M. E., & Cara, A. (2004). Novel integrase-defective lentiviral episomal vectors for gene transfer. Human Gene Therapy, 15(4), 361–372. https://doi.org/10.1089/104303404322959515

Vockley, C. M., D’Ippolito, A. M., McDowell, I. C., Majoros, W. H., Safi, A., Song, L., Crawford, G. E., & Reddy, T. E. (2016). Direct GR Binding Sites Potentiate Clusters of TF Binding across the Human Genome. Cell, 166(5), 1269-1281.e19. https://doi.org/10.1016/j.cell.2016.07.049

Wang, X., He, L., Goggin, S. M., Saadat, A., Wang, L., Sinnott-Armstrong, N., Claussnitzer, M., & Kellis, M. (2018). High-resolution genome-wide functional dissection of transcriptional regulatory regions and nucleotides in human. Nature Communications, 9(1). https://doi.org/10.1038/s41467-018-07746-1

Wanisch, K., & Yáñez-Muñoz, R. J. (2009). Integration-deficient lentiviral vectors: A slow coming of age. In Molecular Therapy (Vol. 17, Issue 8, pp. 1316–1332). Mol Ther. https://doi.org/10.1038/mt.2009.122

Yu, S. F., von Ruden, T., Kantoff, P. W., Garber, C., Seiberg, M., Rüther, U., Anderson, W. F., Wagner, E. F., & Gilboa, E. (1986). Self-inactivating retroviral vectors designed for transfer of whole genes into mammalian cells. Proceedings of the National Academy of Sciences of the United States of America, 83(10), 3194–3198. https://doi.org/10.1073/pnas.83.10.3194

