## Supplemental Figure 1 for "Adaptation of STARR-seq method to be used with 3^rd^ generation integrase-deficient promoterless lentiviral vectors"

**Figure S1 | Map and full sequence of Lenti-STARR-seq plasmid with minCMV promoter.**

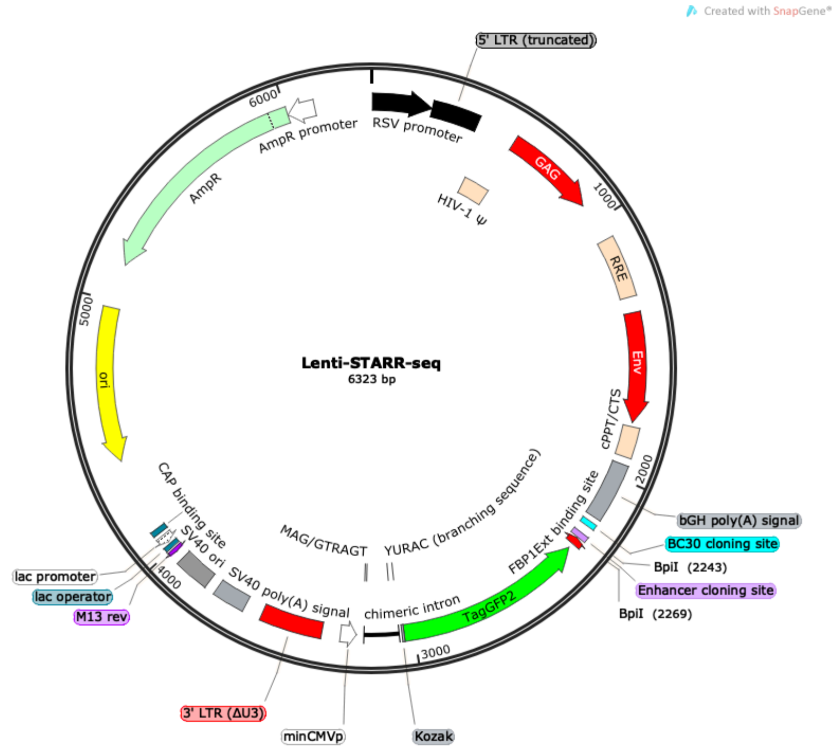


LOCUS Exported 6323 bp ds-DNA circular SYN 24-JUN-2020
DEFINITION synthetic circular DNA
ACCESSION
VERSION
KEYWORDS Lenti-STARR-seq
SOURCE synthetic DNA construct
ORGANISM synthetic DNA construct
REFERENCE 1 (bases 1 to 6323)
AUTHORS CellAge
TITLE Direct Submission
JOURNAL Exported 24 Jun 2020 from SnapGene 2.7.3 http://www.snapgene.com
FEATURES Location/Qualifiers
source 1..6323
/organism="synthetic DNA construct"
/mol_type="other DNA"
source 3053..3195
/organism="synthetic DNA construct"
/mol_type="other DNA"
promoter 6..232
/note="RSV promoter"
/note="Rous sarcoma virus enhancer/promoter"
LTR 233..413
/note="5' LTR (truncated)"
/note="truncated 5' long terminal repeat (LTR) from HIV-1"
misc_feature 460..585
/note="HIV-1 Psi"
/note="packaging signal of human immunodeficiency virus
type 1"

misc_feature 569..922
/note="GAG"
/note="SHould not be here in 2/3rd gen vector?"
misc_feature 1078..1311
/note="RRE"
/note="The Rev response element (RRE) of HIV-1 allows for
Rev-dependent mRNA export from the nucleus to the
cytoplasm."
misc_feature 1371..1787
/note="Env"
/note="SHould not be here in 2/3rd gen vector?"
misc_feature 1803..1918
/note="cPPT/CTS"
/note="central polypurine tract and central termination
sequence of HIV-1 (lacking the first T)"
polyA_signal 1948..2172
/note="bGH poly(A) signal"
/note="bovine growth hormone polyadenylation signal"
misc_feature 2196..2214
/note="BC30 cloning site"
misc_feature 2248..2269
/note="Enhancer cloning site"
misc_feature complement(2274..2301)
/note="FBP1Ext binding site"
/note="This is for primer cloning?"
CDS complement(2322..3038)
/codon_start=1
/product="monomeric green fluorescent protein, also known
as mTagGFP"
/note="TagGFP2"
/note="mammalian codon-optimized"
/translation="MSGGEELFAGIVPVLIELDGDVHGHKFSVRGEGEGDADYGKLEIK

FICTTGKLPVPWPTLVTTLCYGIQCFARYPEHMKMNDFFKSAMPEGYIQERTIQFQDDG

KYKTRGEVKFEGDTLVNRIELKGKDFKEDGNILGHKLEYSFNSHNVYIRPDKANNGLEA

NFKTRHNIEGGGVQLADHYQTNVPLGDGPVLIPINHYLSTQTKISKDRNEARDHMVLLE
SFSACCHTHGMDELYR"
misc_feature 3039..3046
/note="Kozak"
misc_feature complement(3056..3059)
/note="CAG/G"
intron 3057..3189
/label=chimeric intron
/note="chimeric intron"
/note="chimera between introns from human beta-globin and
immunoglobulin heavy chain genes"
misc_feature complement(3082..3086)
/note="YURAC (branching sequence)"
misc_feature complement(3184..3192)
/note="MAG/GTRAGT"
misc_feature complement(3219..3277)
/note="minCMVp"
LTR 3349..3582

/note="3' LTR (Delta-U3)"
/note="self-inactivating 3' long terminal repeat (LTR) from
HIV-1"
polyA_signal 3654..3775
/note="SV40 poly(A) signal"
/note="SV40 polyadenylation signal"
rep_origin 3815..3950
/note="SV40 ori"
/note="SV40 origin of replication

Another SV40??"
primer_bind complement(3983..3999)
/note="M13 rev"
/note="common sequencing primer, one of multiple similar
variants"
protein_bind 4007..4023
/bound_moiety="lac repressor encoded by lacI"
/note="lac operator"
/note="The lac repressor binds to the lac operator to
inhibit transcription in E. coli. This inhibition can be
relieved by adding lactose or
isopropyl-beta-D-thiogalactopyranoside (IPTG)."
promoter complement(4031..4061)
/note="lac promoter"
/note="promoter for the E. coli lac operon"
protein_bind 4076..4097
/bound_moiety="E. coli catabolite activator protein"
/note="CAP binding site"
/note="CAP binding activates transcription in the presence
of cAMP."
rep_origin complement(4385..4973)
/direction=LEFT
/note="ori"
/note="high-copy-number ColE1/pMB1/pBR322/pUC origin of
replication"
CDS complement(5144..6004)
/codon_start=1
/gene="bla"
/product="beta-lactamase"
/note="AmpR"
/note="confers resistance to ampicillin, carbenicillin, and
related antibiotics"
/translation="MSIQHFRVALIPFFAAFCLPVFAHPETLVKVKDAEDQLGARVGYI

ELDLNSGKILESFRPEERFPMMSTFKVLLCGAVLSRIDAGQEQLGRRIHYSQNDLVEYS

PVTEKHLTDGMTVRELCSAAITMSDNTAANLLLTTIGGPKELTAFLHNMGDHVTRLDRW

EPELNEAIPNDERDTTMPVAMATTLRKLLTGELLTLASRQQLIDWMEADKVAGPLLRSA

LPAGWFIADKSGAGERGSRGIIAALGPDGKPSRIVVIYTTGSQATMDERNRQIAEIGAS
LIKHW"
promoter complement(6005..6109)
/gene="bla"
/note="AmpR promoter"

ORIGIN
1 acgcgtgtag tcttatgcaa tactcttgta gtcttgcaac atggtaacga tgagttagca
61 acatgcctta caaggagaga aaaagcaccg tgcatgccga ttggtggaag taaggtggta
121 cgatcgtgcc ttattaggaa ggcaacagac gggtctgaca tggattggac gaaccactga
181 attgccgcat tgcagagata ttgtatttaa gtgcctagct cgatacataa acgggtctct
241 ctggttagac cagatctgag cctgggagct ctctggctaa ctagggaacc cactgcttaa
301 gcctcaataa agcttgcctt gagtgcttca agtagtgtgt gcccgtctgt tgtgtgactc
361 tggtaactag agatccctca gacccttttta gtcagtgtgg aaaatctcta gcagtggcgc
421 ccgaacaggg acttgaaagc gaaagggaaa ccagaggagc tctctcgacg caggactcgg
481 cttgctgaag cgcgcacggc aagaggcgag gggcggcgac tggtgagtac gccaaaaatt
541 ttgactagcg gaggctagaa ggagagagat gggtgcgaga gcgtcagtat taagcggggg
601 agaattagat cgcgatggga aaaaattcgg ttaaggccag ggggaaagaa aaaatataaa
661 ttaaaacata tagtatgggc aagcagggag ctagaacgat tcgcagttaa tcctggcctg
721 ttagaaacat cagaaggctg tagacaaata ctgggacagc tacaaccatc ccttcagaca
781 ggatcagaag aacttagatc attatataat acagtagcaa ccctctattg tgtgcatcaa
841 aggatagaga taaaagacac caaggaagct ttagacaaga tagaggaaga gcaaaacaaa
901 agtaagacca ccgcacagca agcggccact gatcttcaga cctggaggag gagatatgag
961 ggacaattgg agaagtgaat tatataaata taaagtagta aaaattgaac cattaggagt
1021 agcacccacc aaggcaaaga gaagagtggt gcagagagaa aaaagagcag tgggaatagg
1081 agctttgttc cttgggttct tgggagcagc aggaagcact atgggcgcag cgtcaatgac
1141 gctgacggta caggccagac aattattgtc tggtatagtg cagcagcaga acaatttgct
1201 gagggctatt gaggcgcaac agcatctgtt gcaactcaca gtctggggca tcaagcagct
1261 ccaggcaaga atcctggctg tggaaagata cctaaaggat caacagctcc tggggatttg
1321 gggttgctct ggaaaactca tttgcaccac tgctgtgcct tggaatgcta gttggagtaa
1381 taaatctctg gaacagattt ggaatcacac gacctggatg gagtgggaca gagaaattaa
1441 caattacaca agcttaatac actccttaat tgaagaatcg caaaaccagc aagaaaagaa
1501 tgaacaagaa ttattggaat tagataaatg ggcaagtttg tggaattggt ttaacataac
1561 aaattggctg tggtatataa aattattcat aatgatagta ggaggcttgg taggtttaag
1621 aatagttttt gctgtacttt ctatagtgaa tagagttagg cagggatatt caccattatc
1681 gtttcagacc cacctcccaa ccccgagggg acccgacagg cccgaaggaa tagaagaaga
1741 aggtggagag agagacagag acagatccat tcgattagtg aacggatctc gacggtatcg
1801 aatttaaaag aaaagggggg attggggggt acagtgcagg ggaaagaata gtagacataa
1861 tagcaacaga catacaaact aaagaattac aaaaacaaat tacaaaaatt caaaatttat
1921 cgattctgaa gctttccgcc tcagaagcca tagagcccac cgcatcccca gcatgcctgc
1981 tattgacttc ccaatcctcc cccttgctgt cctgccccac cccacccccc agaatagaat
2041 gacacctact cagacaatgc gatgcaattt cctcatttta ttaggaaagg acagtgggag
2101 tggcaccttc cagggtcaag gaaggcacgg gggaggggca aacaacagat ggctggcaac
2161 tagaaggcac agtcgaggca ccggtgaatt cacgatgaga cgaaaaaacg tctcgtgcgg
2221 gaggaggtga ggctacctga gcattccgtg tcttcaaaaa agaagactca tcgtgcgtgc
2281 gttgcgttcg gtggtcggtc ggaattcgtc gacaatctag ctcacctgta cagctcgtcc
2341 atgccgtggg tgtggcagca ggcgctgaag gactccagga gcaccatgtg gtcgcgggcc
2401 tcgttgcggt ccttgctgat cttggtctga gtgctcaggt agtggttgat ggggatcagc
2461 acggggccgt cgcccagggg cacgttggtc tggtagtggt cggccagctg cacgccgccg
2521 ccctcgatgt tgtggcgggt cttgaagtta gcctccaggc cgttgttggc cttgtcgggg
2581 cggatgtaca cgttgtggct gttgaagctg tactccagct tgtggcccag gatgttgccg
2641 tcctccttga agtccttgcc cttcagctcg atgcggttca ccagggtgtc gccctcgaac
2701 ttcacctcgc cgcgggtctt gtacttgccg tcgtcctgga actggatggt gcgctcctgg
2761 atgtagccct cgggcatggc gctcttgaag aagtcgttca tcttcatgtg ctcggggtag
2821 cgggcgaagc actggatgcc gtagcagagg gtggtcacca gggtgggcca gggcacgggc
2881 agcttgccgg tggtgcagat gaacttgatc tccagcttgc cgtagtcggc gtcgccctcg
2941 ccctcgccgc gcacgctgaa cttgtggccg tgcacgtcgc cgtccagctc gatcagcacg
3001 ggcacgatgc cggcgaacag ctcctcgccc ccgctcatgg tggcgtctag acacacctgt
3061 ggagagaaag gcaaagtgga tgtcagtaag accaataggt gcctatcaga aacgcaagag
3121 tcttgtctgt ctcgacaagc ccagtttcta ttggtctcct taaacctgtc ttgtaacctt
3181 gatacttacc tgccctagac tcgagaatcg acgaattaga tctgacggtt cactaaacga

3241 gctctgctta tatagacctc ccaccgtaca cgcctacggt acctttaaga ccaatgactt
3301 acaaggcagc tgtagatctt agccactttt taaaagaaaa ggggggactg gaagggctaa
3361 ttcactccca acgaagataa gatctgcttt ttgcttgtac tgggtctctc tggttagacc
3421 agatctgagc ctgggagctc tctggctaac tagggaaccc actgcttaag cctcaataaa
3481 gcttgccttg agtgcttcaa gtagtgtgtg cccgtctgtt gtgtgactct ggtaactaga
3541 gatccctcag acccttttag tcagtgtgga aaatctctag cagtagtagt tcatgtcatc
3601 ttattattca gtatttataa cttgcaaaga aatgaatatc agagagtgag aggaacttgt
3661 ttattgcagc ttataatggt tacaaataaa gcaatagcat cacaaatttc acaaataaag
3721 catttttttc actgcattct agttgtggtt tgtccaaact catcaatgta tcttatcatg
3781 tctggctcta gctatcccgc ccctaactcc gcccatcccg cccctaactc cgcccagttc
3841 cgcccattct ccgccccatg gctgactaat tttttttatt tatgcagagg ccgaggccgc
3901 ctcggcctct gagctattcc agaagtagtg aggaggcttt tttggaggcc tagacttttg
3961 cagagaccaa attcgtaatc atgtcatagc tgtttcctgt gtgaaattgt tatccgctca
4021 caattccaca caacatacga gccggaagca taaagtgtaa agcctggggt gcctaatgag
4081 tgagctaact cacattaatt gcgttgcgct cactgcccgc tttccagtcg ggaaacctgt
4141 cgtgccagct gcattaatga atcggccaac gcgcggggag aggcggtttg cgtattgggc
4201 gctcttccgc ttcctcgctc actgactcgc tgcgctcggt cgttcggctg cggcgagcgg
4261 tatcagctca ctcaaaggcg gtaatacggt tatccacaga atcaggggat aacgcaggaa
4321 agaacatgtg agcaaaaggc cagcaaaagg ccaggaaccg taaaaaggcc gcgttgctgg
4381 cgtttttcca taggctccgc ccccctgacg agcatcacaa aaatcgacgc tcaagtcaga
4441 ggtggcgaaa cccgacagga ctataaagat accaggcgtt tccccctgga agctccctcg
4501 tgcgctctcc tgttccgacc ctgccgctta ccggatacct gtccgccttt ctcccttcgg
4561 gaagcgtggc gctttctcat agctcacgct gtaggtatct cagttcggtg taggtcgttc
4621 gctccaagct gggctgtgtg cacgaacccc ccgttcagcc cgaccgctgc gccttatccg
4681 gtaactatcg tcttgagtcc aacccggtaa gacacgactt atcgccactg gcagcagcca
4741 ctggtaacag gattagcaga gcgaggtatg taggcggtgc tacagagttc ttgaagtggt
4801 ggcctaacta cggctacact agaagaacag tatttggtat ctgcgctctg ctgaagccag
4861 ttaccttcgg aaaaagagtt ggtagctctt gatccggcaa acaaaccacc gctggtagcg
4921 gtggtttttt tgtttgcaag cagcagatta cgcgcagaaa aaaaggatct caagaagatc
4981 ctttgatctt ttctacgggg tctgacgctc agtggaacga aaactcacgt taagggattt
5041 tggtcatgag attatcaaaa aggatcttca cctagatcct tttaaattaa aaatgaagtt
5101 ttaaatcaat ctaaagtata tatgagtaaa cttggtctga cagttaccaa tgcttaatca
5161 gtgaggcacc tatctcagcg atctgtctat ttcgttcatc catagttgcc tgactccccg
5221 tcgtgtagat aactacgata cgggagggct taccatctgg ccccagtgct gcaatgatac
5281 cgcgagaccc acgctcaccg gctccagatt tatcagcaat aaaccagcca gccggaaggg
5341 ccgagcgcag aagtggtcct gcaactttat ccgcctccat ccagtctatt aattgttgcc
5401 gggaagctag agtaagtagt tcgccagtta atagtttgcg caacgttgtt gccattgcta
5461 caggcatcgt ggtgtcacgc tcgtcgtttg gtatggcttc attcagctcc ggttcccaac
5521 gatcaaggcg agttacatga tcccccatgt tgtgcaaaaa agcggttagc tccttcggtc
5581 ctccgatcgt tgtcagaagt aagttggccg cagtgttatc actcatggtt atggcagcac
5641 tgcataattc tcttactgtc atgccatccg taagatgctt ttctgtgact ggtgagtact
5701 caaccaagtc attctgagaa tagtgtatgc ggcgaccgag ttgctcttgc ccggcgtcaa
5761 tacgggataa taccgcgcca catagcagaa ctttaaaagt gctcatcatt ggaaaacgtt
5821 cttcggggcg aaaactctca aggatcttac cgctgttgag atccagttcg atgtaaccca
5881 ctcgtgcacc caactgatct tcagcatctt ttactttcac cagcgtttct gggtgagcaa
5941 aaacaggaag gcaaaatgcc gcaaaaaagg gaataagggc gacacggaaa tgttgaatac
6001 tcatactctt cctttttcaa tattattgaa gcatttatca gggttattgt ctcatgagcg
6061 gatacatatt tgaatgtatt tagaaaaata aacaaatagg ggttccgcgc acatttcccc
6121 gaaaagtgcc acctgacgtc aacggatcgg gagatcaact ggataactca agctaaccaa
6181 aatcatccca aacttcccac cccatacct attaccactg ccaattacct gtggtttcat
6241 ttactctaaa cctgtgattc ctctgaatta ttttcatttt aaagaaattg tatttgttaa
6301 atatgtacta caaacttagt agt
//
